# Sleep spindles mediate hippocampal-neocortical coupling during sharp-wave ripples

**DOI:** 10.1101/712463

**Authors:** Hong-Viet. V. Ngo, Juergen Fell, Bernhard P. Staresina

## Abstract

Sleep is pivotal for the consolidation of memories [1]. According to two-stage accounts, experiences are temporarily stored in the hippocampus and gradually translocated to neocortical sites during non-rapid-eye-movement (NREM) sleep [2,3]. Mechanistically, information transfer is thought to rely on interactions between thalamocortical spindles and hippocampal ripples. In particular, spindles may open precisely-timed communication channels, across which reactivation patterns may travel between the hippocampus and cortical target sites when ripples occur. To test this hypothesis, we first derived time-frequency representations (TFRs) in hippocampus (HIPP) and at scalp electrode Cz (neocortex, NC) time-locked to individual hippocampal ripple events. Compared to matched ripple-free intervals, results revealed a concurrent increase in spindle power both in HIPP and NC. As revealed by coherence analysis, hippocampal-neocortical coupling was indeed enhanced in the spindle band around ripples. Finally, we examined the directionality of spindle coupling and observed a strong driving effect from NC to HIPP. Specifically, ∼250 ms prior to the HIPP ripple, NC spindles emerge and entrain HIPP spindles. Both regions then remain synchronised until ∼500 ms after the ripple. Consistent with recent rodent work, these findings suggest that active consolidation is initiated by neocortex and draws on neocortical-hippocampal-neocortical reactivation loops [4], with a role of sleep spindles in mediating this process.

## Introduction

Information transfer during sleep is thought to rely on systematic interactions of the cardinal NREM sleep rhythms [5]: First, the cortical <1 Hz slow oscillation (SO) opens windows of neuronal excitability and inhibition (up- and down-states, respectively) in cortical and subcortical regions [6,7]. Triggered by SOs, the thalamus generates sleep spindles, i.e., transient (0.5-2 s) oscillatory activity between 12-16 Hz, via thalamo-cortical loops [8]. Nested in SO up-states, spindles gate Ca^2+^ influx into dendrites and promote synaptic plasticity [9–11]. Importantly, spindles have also been shown to group hippocampal sharp-wave ripples (SW-Rs) [12–14]. SW-Rs are transient events emerging from recurrent interactions between the CA1 and CA3 subfields and consist of a brief 80-200-Hz ripple-burst superimposed on a ∼3-Hz sharp wave [14–16]. Ripples have been linked to the reactivation of cell assemblies engaged during previous encoding [17–19], and experimental suppression of ripples leads to an impairment of memory performance [20,21]. Given that spindles group SW-Rs and induce neural plasticity, they appear ideally-suited to facilitate memory consolidation mechanistically. That is, spindles might synchronise the hippocampus (sender) and cortical target sites (receiver) during reactivation events and thereby induce synaptic changes in neocortex for long-term storage [22,23]. However, whether and how spindles mediate the hippocampal-neocortical dialogue still remains poorly understood. We thus examined whole-night sleep recordings from neocortex and hippocampus in 14 pre-surgical epilepsy patients (Figure 1). In a first step, we algorithmically detected spindles in neocortex (NC; scalp electrode Cz) and posterior hippocampus (HIPP; [14]) as well as hippocampal sharp-wave ripples (SW-Rs). Across participants, a total of 17,174 NC spindles, 14,748 HIPP spindles and 9,038 HIPP SW-Rs were identified (Figure 1). For participant-specific sleep characteristics, event numbers and densities (events per minute), see Table 1.

**Table 1.** Sleep architecture and properties of sleep spindles and sharp-wave ripples. Mean ± s.e.m. proportion of sleep stages S1, S2, slow wave sleep (SWS) and rapid eye movement (REM) sleep relative to the total time spent asleep. Mean (± s.e.m.) count and density (events per min) of offline detected spindles and sharp-wave ripples (SW-Rs) across participants.

|  | Mean | s.e.m. |
| --- | --- | --- |
| S1 (%) | 22.8 | 4.6 |
| S2 (%) | 44.0 | 3.3 |
| SWS (%) | 16.4 | 2.6 |
| REM (%) | 16.8 | 2.2 |
| Total sleep time (min) | 427.6 | 29.0 |
| Spindle count |  |  |
| NC | 1226.7 | 125.9 |
| HIPP | 1053.4 | 150.6 |
| Spindle density (per min) |  |  |
| NC | 5.2 | 0.3 |
| HIPP | 4.9 | 0.4 |
| SW-R count | 645.6 | 121.7 |
| SW-R density (per min) | 2.9 | 0.4 |

**Figure 1.**
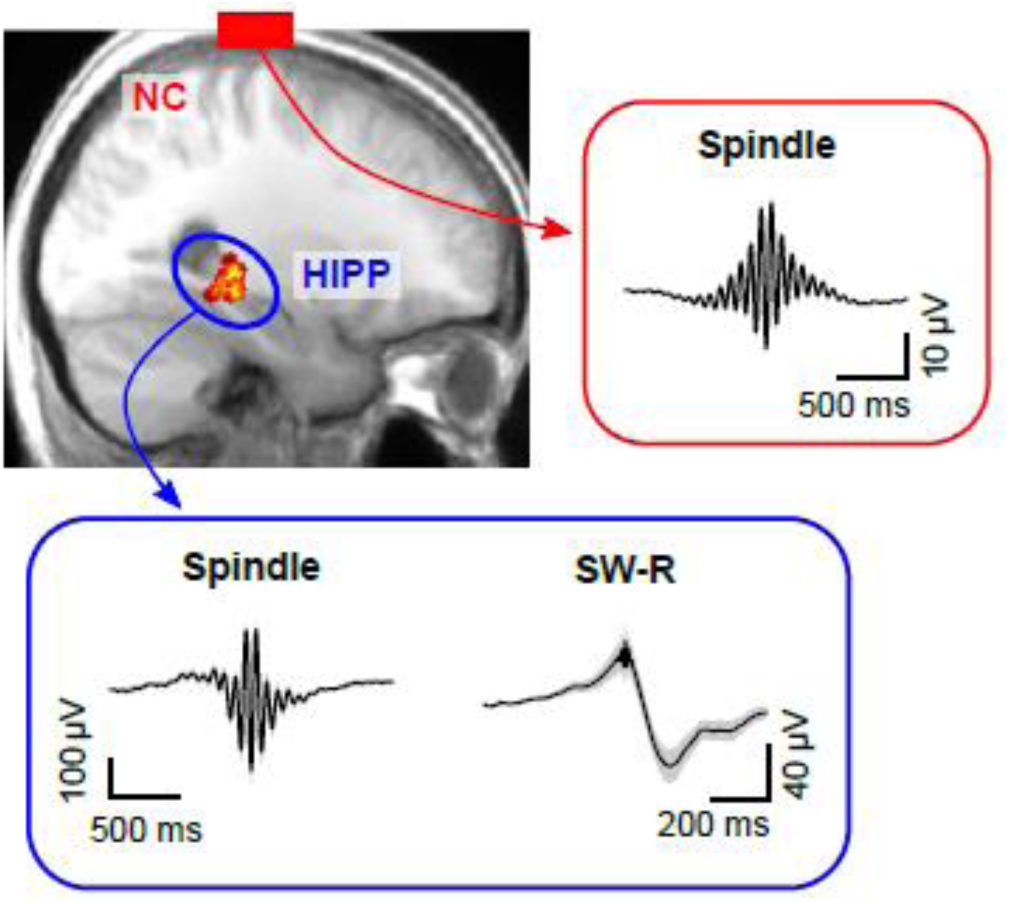
Cortical sleep spindles and hippocampal spindles and SW-Rs. *Top left*: heat map illustrating the position of individual contacts across all patients within the hippocampus overlaid on a sagittal slice of the mean structural MRI. *Right* and *bottom* insets show the grand average (± s.e.m.) spindle and sharp-wave ripple waveforms across all patients within the neocortical (red, NC) and hippocampus (blue, HIPP).

## Results

### Cortical and hippocampal spindles around SW-Rs

To examine whether SW-Rs co-occur not only with spindles in HIPP [14] but also in NC, we first derived time-frequency representations (TFRs) time-locked to discrete HIPP SW-Rs (Figure 2). Results were statistically compared to TFRs obtained from surrogate events, i.e., matching ripple-free intervals randomly drawn from NREM sleep (see Material and Methods). In HIPP, we found an extended cluster of significant power increases encompassing three distinct components (P = 0.001, Figure 2A): First, an increase in SO power (< 1 Hz), ranging from ∼-1.0 to 0.5 s relative to SW-Rs. Second, an increase in spindle power (12-18 Hz), onsetting prior to and peaking at ∼300 ms after SW-Rs. Third, a more widespread frequency cluster with a maximum at 3 Hz, reflecting the sharp-wave component [14,24,25]. Critically, when locking the TFR in NC to SW-Rs in HIPP (Figure 2B), we again observed significant power increases compared to surrogate data. A significant cluster emerged from 11 to 16 Hz (P = 0.001), again arising before and peaking after SW-Rs. The concurrent spindle power increase in both regions (Figure 2C), emerging prior to the HIPP SW-R and peaking shortly thereafter, might indicate a role of spindles in mediating cortical-hippocampal communication (see next section).

**Figure 2.**
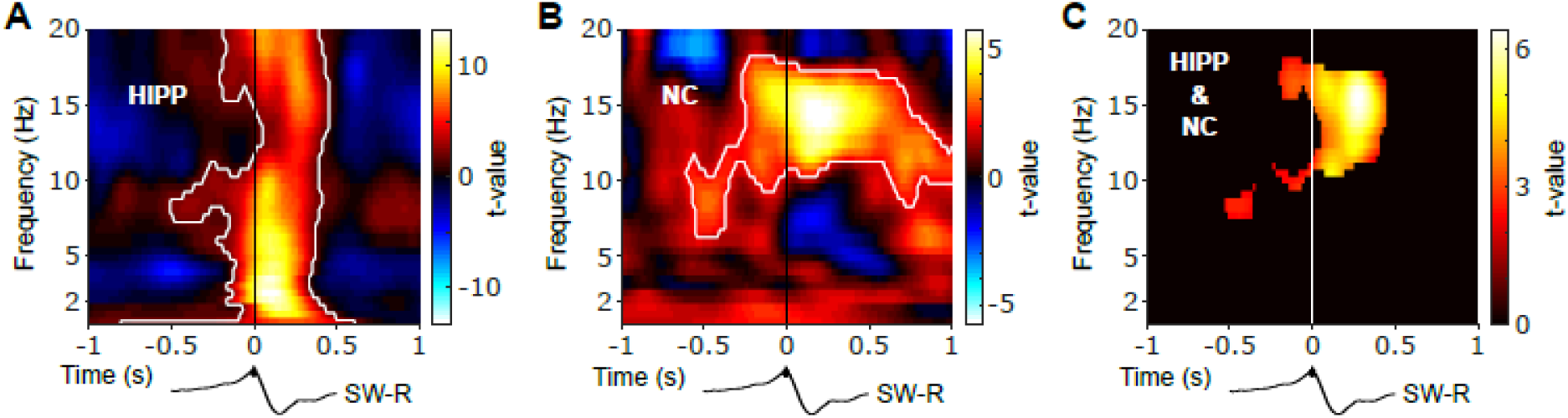
Neocortical and hippocampal spindle activity coincides during SW-Rs. Statistical maps (t-values) contrasting ripple-locked and surrogate TFRs within HIPP (A) and NC (B). Hot colors indicate power increases around SW-Rs, whereas blue colours indicate relative power decreases. White contours mark significant clusters obtained from a cluster-based permutation procedure. (C) Significance mask derived from the overlap of significant clusters between NC and HIPP. Color represents the mean t-value from the corresponding statistical masks. Black traces below illustrate the timing of power changes relative to SW-Rs, which particularly highlights the power increase in HIPP at ∼3 Hz reflecting the sharp wave component in (A).

Of note, theoretical accounts and empirical work has also implicated SOs/slow-waves in the hippocampal-neocortical dialogue [13,26]. While the ripple-locked HIPP-TFR showed a significant increase at ∼1 Hz relative to surrogates (Figure 2A), no such effect was seen in the NC-TFR (Figure 2B). However, this might result from our surrogate procedure, which extracts matched NREM epochs without HIPP SW-Rs (see Material and Methods). That is, the prevalence of neocortical SOs throughout NREM sleep (thus also in surrogate epochs) might obscure SOs aligned to SW-Rs. Indeed, when comparing ripple-locked TFRs with a −2 s to −1.5 s pre-ripple interval, a significant power increase was seen in both the spindle and SO band in both HIPP and NC (Figure S1). That said, comparison with matched ripple-free surrogates has the advantage of avoiding the arbitrary choice of a pre-event baseline interval. In any case, the fact that spindles emerged in the ripple-locked TFRs of both regions relative to NREM surrogates emphasises that spindles cluster around HIPP ripples above and beyond background spindle densities.

### Cortico-hippocampal spindle coupling around SW-Rs

Next, we asked whether the cross-regional co-activation in the spindle band around HIPP SW-Rs may indeed reflect an increase in functional coupling between NC and HIPP. To this end, we calculated ripple-locked spectral coherence between NC and HIPP. Based on the concurrent power increases shown in Figure 2, we focused on the 12-16 spindle range and determined coherence in a symmetric 500 ms around SW-Rs. Note though that the results remain unchanged when reducing this window to 400 ms or expanding it up to 800 ms (for extended time-frequency resolved coherence analysis, see Figure S2). As shown in Figure 3A, spindle-band coherence between NC and HIPP was indeed significantly increased compared to surrogates (z-value = 2.118, P = 0.034). Examination of time-resolved 12-16 Hz coherence confirmed a significant increase starting ∼100 ms before the SW-R and reaching its maximum shortly after the SW-R (Figure 3B). Identical results were obtained using amplitude- or phase-based connectivity measures (Figure S3), i.e. orthogonalized power correlation [27] and phase-locking value [28], which validated the robustness of our coherence-based approach and - more importantly - ruled out any influence of volume conduction or spurious correlations due to a common referencing scheme. Altogether, these findings corroborate the notion that sleep spindles co-occurring in NC and HIPP reflect an increase in cortico-hippocampal communication.

**Figure 3.**
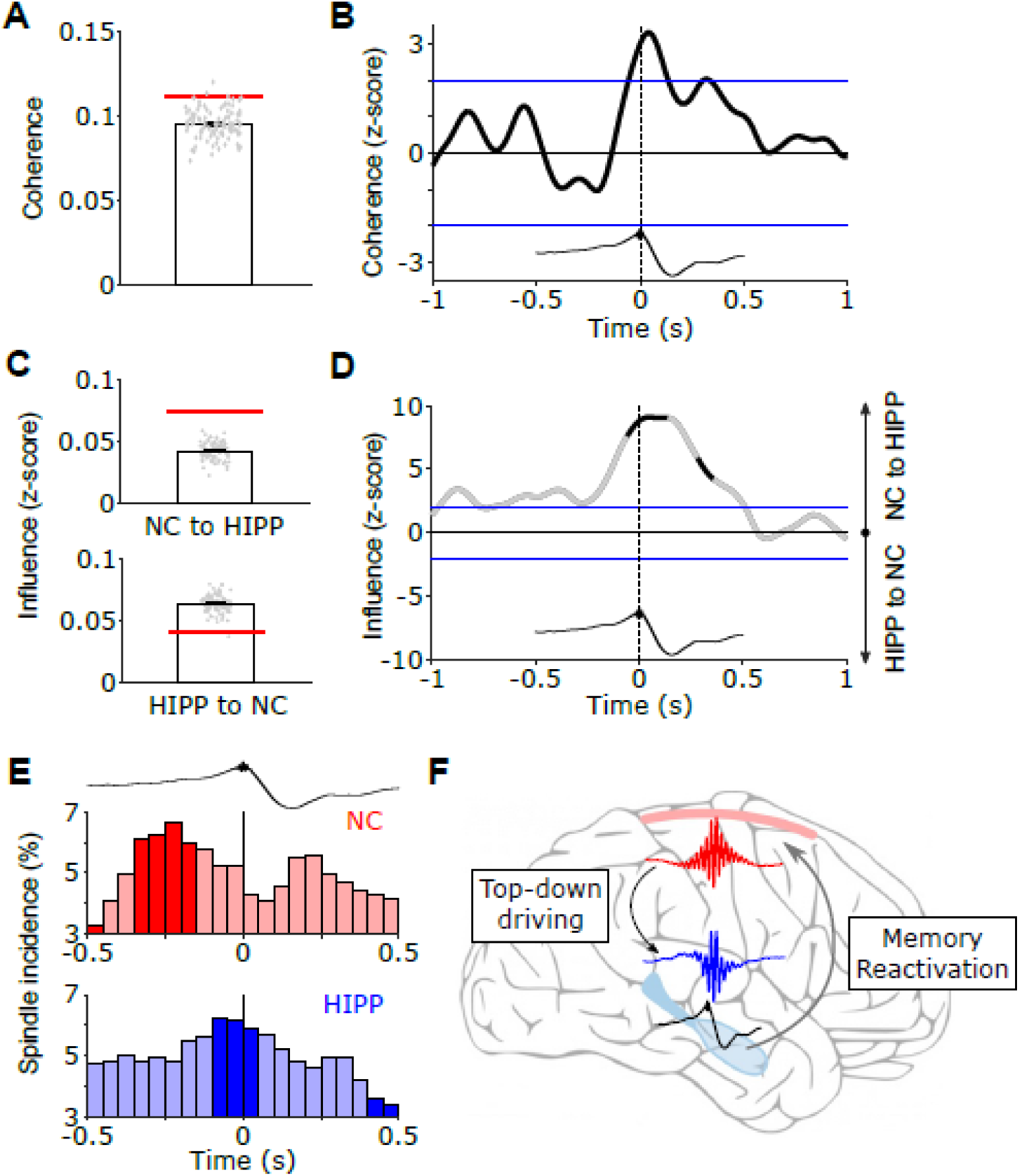
Top-down-initiated cortico-hippocampal communication via spindles. (A) HIPP-NC coherence. The red line depicts the observed coherence between HIPP and NC in the 12-16 Hz frequency range centred from −0.25 to +0.25 s around HIPP SW-Rs. Bar graph represents mean coherence ± s.e.m. for ripple-free surrogate events. Grey circles illustrate individual values for all 100 surrogates. *p < 0.05 for comparison between ripple and surrogate data (z-score). (B) Time-resolved HIPP-NC coherence for the 12-16 Hz spindle range transformed into a z-score with respect to the surrogate events. Blue lines indicate standard significance thresholds (z = 1.96). Time 0 denotes HIPP SW-Rs. (C) Partial directed coherence (PDC). Red lines represent top-down directionality from NC to HIPP (*top*) or bottom-up directionality from HIPP to NC (*bottom*) in the 12-16 Hz frequency range centred from −0.25 to +0.25 s around HIPP SW-Rs. Bar graphs represent mean PDC ± s.e.m. for corresponding surrogates. Grey circles illustrate distribution of values across all 100 sets of surrogates. (D) Time course of difference between z-transformed top-down and bottom-up influence in the 12-16 Hz spindle range. Positive values signify a cortical spindle influence on HIPP and vice versa for negative values. Black-coloured sections correspond to time intervals with significant spectral coherence shown in Figure 3B. Blue lines mark standard significance thresholds (z = 1.96). (E) Peri-event histograms of spindle onsets in NC (*top*) and HIPP (*bottom*) within a ± 0.5 s time window around SW-Rs (time = 0 s, top trace). Dark coloured bars indicate significant time bins, resulting from comparison with ripple-free surrogate events (z > 1.96). (F) Schematic illustrating the spindle-mediated cortico-hippocampal dialogue around SW-Rs: First, NC-spindles drive HIPP-spindles in a top-down fashion, which in turn coordinate the occurrence of SW-Rs on a fine temporal scale. SW-Rs are linked to the reactivation of relevant memory traces, assumed to be distributed to neocortical sites for long-term storage.

### Cortical spindles drive hippocampal spindles prior to SW-Rs

Lastly, we asked whether this cross-regional spindle interaction is directional, i.e., do cortical spindles influence hippocampal spindles or vice versa? As a measure of directionality, we used partial directed coherence (PDC). Extending the concept of coherence, i.e. mutual synchronous activity, PDC disentangles how the current state of a target region is influenced by the past of other regions or of the target region itself [29]. Calculating 12-16 Hz PDC again from −0.25 to +0.25 s around SW-Rs (for extended time-frequency resolved PDC analysis see Figure S4) revealed an increase in directional influence primarily from the neocortex to the hippocampus in comparison to surrogates (Figure 3C, z = 4.763, P < 0.001). The inverse directionality (HIPP -> NC) was diminished around SW-Rs in comparison to surrogates (z = −3.170, P = 0.002). The direct comparison of the directional influence between cortical and hippocampal spindles revealed an almost two-fold influence of neocortical spindles on hippocampal spindles than vice versa. Again, inspecting the temporal dynamics of PDC within the spindle range corroborated that the top-down influence sets in before the occurrence of HIPP SW-Rs (Figure 3D). To further examine the origin of the directional influence of NC on HIPP spindles, we extracted the onset latencies of discrete spindles in both regions with respect to HIPP SW-Rs. Note that while the TFR analysis shown in Figure 2 highlighted significant increases in amplitude, this analysis is particularly geared towards detecting spindle onsets. As shown in Figure 3E, histograms of spindle onsets revealed a maximum (significantly increasing compared to surrogates) in both regions before the SW-Rs. Importantly though, this maximum occurred from −250 to −200 ms prior to the hippocampal ripple in NC and from −100 to −50 ms prior to the ripple in HIPP. Together, these results suggest a driving role of NC spindles on HIPP spindles prior to the SW-R.

## Discussion

Memory consolidation is thought to rely on an intricate interplay between SOs, sleep spindles and SW-Rs. While the link between each of these sleep signatures and effective consolidation has been established across species [20,21,26,30–33], less is known about their dynamic interactions underlying the purported information transfer from hippocampus to neocortex [5]. In particular, it has been argued that sleep spindles might be pivotal for facilitating hippocampal-neocortical interactions, opening fine-tuned windows of opportunity for cross-regional synchronisation and plasticity [10,11,22,34–36]. In other words, spindles might serve as the mechanistic vehicle to synchronise the hippocampus with neocortical target sites at the time of hippocampal memory reactivation, i.e., around SW-Rs. In rodents, hippocampal ripples have been shown to coincide with spindles in prefrontal cortex [12], and we recently showed that spindles group SW-Rs within the hippocampus [14]. Critically though, whether hippocampus and neocortical sites are indeed synchronised by spindles at the time of SW-Rs has remained open. Here we analysed EEG data acquired during NREM sleep both in hippocampus (HIPP) and at scalp electrode Cz (Figure 1), the latter integrating across fronto-parietal neocortical (NC) networks. Time-locking the continuous EEG to individual HIPP SW-Rs, we first showed that spindle power (∼12-16 Hz) increases not only in HIPP around SW-Rs [14], but also in NC (Figure 2). Next, using coherence analysis, we were able to show an increase in HIPP-NC functional coupling in the spindle band around ripples (Figure 3A and S2). These results are consistent with a role of spindles in mediating the information transfer between HIPP and NC around SW-Rs, i.e. when memory traces are presumably reactivated in HIPP [37,38].

One critical question though is how the hippocampal-neocortical dialogue is initiated in the sleeping brain, i.e. what governs the interplay between spindles and SW-Rs (and SOs). One intuitive scenario is that upon spontaneous SW-Rs in HIPP, sleep spindles – projected from the thalamus to both HIPP and NC – synchronise the two regions. Indeed, rodent recordings have shown that prefrontal cortex neurons fire after HIPP neurons during slow wave sleep [12], and the stronger the hippocampal firing burst, the more likely a spindle event is to be observed in PFC [39]. An alternative scenario, however, is that HIPP-NC interactions are initiated in neocortex, perhaps ensuring that cortical target sites are in a state conducive to plasticity [4]. Rodent recordings from somatosensory cortex and HIPP have shown that the NC spindles indeed trigger HIPP firing and associated ripples, with NC spindles emerging ∼200 ms before HIPP ripples [13]. Moreover, a recent rodent study revealed a cortical-hippocampal-cortical loop, in that engagement of task-relevant NC sites not only followed reactivation of HIPP assemblies, but also preceded it by ∼200 ms [40]. Of note, HIPP SW-Rs were preceded and accompanied by an increase in NC spindle power. Our current results show a remarkable overlap with these rodent studies regarding the timing of driving NC spindles relative to HIPP ripples (∼200 ms; Figure 3B and D). Moreover, we were further able to show a directional influence of NC to HIPP spindles, pinpointing the role of spindles in synchronising NC and HIPP prior to SW-Rs. In particular, as shown in Figure 3D, partial directed coherence (PDC) strongly suggested a top-down influence of NC on HIPP in the spindle band. This top-down influence emerged ∼250 ms prior to SW-Rs and was sustained until ∼500 ms after the ripple. Corroborating the PDC analysis, we show that the prevalence of spindle onsets rises significantly in both HIPP and NC prior to ripples, but does so sooner in NC than in HIPP (Figure 3E). In sum, these results suggest that the spindle-mediated hippocampal-neocortical dialogue is initiated in NC and is sustained for an extended period after the SW-Rs, allowing for the hypothesised transfer of memory traces reactivated during HIPP SW-Rs (Figure 3F).

Interestingly, in the rodent study mentioned above [40], the authors were also able to elicit the NC-HIPP-NC loop experimentally by playing sounds related to wake learning episodes. Specifically, presentation of the sound led to reactivation in relevant NC sites, followed by HIPP SW-Rs and associated reactivation, followed again by reactivation in NC (although the sound was no longer present). This casts an intriguing light on some recent targeted memory reactivation (TMR) studies in humans [41,42]. Using multivariate decoding methods on scalp EEG data, these studies indicated two phases of memory-related reinstatement after auditory cues: An initial peak within the first second and another peak at ∼2 seconds after cue onset, with concomitant increases in spindle power around both peaks [41]. In light of the results of Rothschild et al. (2017) in conjunction with our current findings, one possibility is that hippocampal reactivation occurs between the two cortical spindle/reactivation events. One speculative scenario is that hippocampal reactivation serves to enrich the cortical memory trace with spatio-temporal episodic details [4].

This still leaves open the question how – in the absence of external cues – the relevant cortical sites get activated in the first place. One parsimonious account would be that although thalamo-cortical spindles are broadcast stochastically, activation thresholds in regions involved in previous learning episodes might be lower due to early plasticity processes set in motion during learning [22,23]. By virtue of synaptic links between NC and HIPP (also strengthened during learning), spindle-activated cortical sites in turn activate the corresponding HIPP assemblies. In other words, functional connectivity between particular NC and HIPP assemblies has already been facilitated during learning [43]. Thalamocortical spindles then entrain these assemblies, first in NC and then in HIPP. The spindle-mediated activation increase in HIPP in turn triggers SW-Rs and associated reactivation of HIPP engrams. The still ongoing spindle ensures that HIPP and NC remain in sync during this reactivation, allowing for further plasticity increases in NC, now incorporating mnemonic elements provided by HIPP. A similar scenario has been elegantly formulated by Rothschild (2018), emphasising the role of NC slow-wave up states in triggering hippocampal reactivation. It deserves explicit mention though that we did not employ a pre-sleep learning task and only recorded from scalp electrode Cz as a proxy for neocortical activation. Our proposal for spindle-mediated reactivation of cortical learning sites thus remains speculative. That said, previous work has indeed shown differential SO and spindle topographies as a function of prior learning tasks [44–47]. In any case, we suggest that the observed top-down spindle effects may reflect cortically-informed memory reactivation [4]: neocortical sleep spindles emerging in brain regions involved in encoding and long-term storage drive hippocampal spindles and SW-Rs, with the latter initiating the distribution of relevant information to the corresponding neocortical networks (Figure 3F).

## Acknowledgments

This research was supported by a Wellcome Trust/Royal Society Sir Henry Dale Fellowship to B.P.S. (107672/Z/15/Z) and a grant from the German Research Foundation to J.F. (FE 366/9-1).

## Author Contributions

Conceptualization: B.P.S. and J.F.; Analysis: H.V.N.; Writing: H.V.N., J.F., B.P.S.

## Material and Methods

### Patients

EEG data from 14 patients (35.4 ± 3.0 years of age, seven females) suffering from pharmacoresistant epilepsy were analysed, which were recorded at the Department of Epileptology, University of Bonn. Intracranial depth electrodes for presurgical evaluation of seizure onset zones were implanted stereotactically, either via the occipital lobe along the longitudinal axis of the hippocampus or laterally via the temporal lobe. Implantations of depth electrodes were bilateral and only electrodes from the non-pathological hemisphere (according to clinical monitoring) were analysed. Informed consent was obtained from all patients and the study was approved by the ethics committee of the Medical Faculty of the University of Bonn.

### Methods

#### EEG recordings and pre-processing

Depth EEG recordings were referenced to linked mastoids and acquired with a sampling rate of 1 kHz (bandpass filter: 0.01 Hz (6 dB per octave) to 300 Hz (12 dB per octave)). For the sleep recordings, additional electrodes were placed on participants’ scalps at positions Cz, C3, C4 and Oz according to the 10-20 system. Electro-ocular activity (EOG) was recorded at the outer canthi of both eyes and submental electromyographic activity (EMG) was acquired with electrodes attached to the chin. Electrode impedances were all below 5 kΩ.

Sleep stages were determined visually using scalp EEG, EOG, and EMG recordings for consecutive 20-s epochs according to standard criteria [48]. For each night, the proportion of sleep stages S1, S2, SWS (i.e., S3 and S4) and REM sleep were calculated relative to the total time spent asleep (Table 1). For pre-processing, an automated algorithm was applied to identify three different types of artifacts separately for each sleep stage: First, on a 0.3 to 150 Hz band-pass filtered signal, amplitude-based artifacts were scored as values exceeding ± 750 µV. Next we identified gradient artifacts, i.e. strong deflections in the signal caused in particular by interictal spikes. Based on the 0.3 to 150 Hz band-pass filtered signal we first calculated for each time-point the difference in amplitude to the next time-point. This difference signal was then used to derive an individual threshold determined by its median ± 6 * interquartile range across all time points. Accordingly, whenever the difference signal exceed this threshold a gradient artifact was scored. Finally, to identity high-frequency bursts emerging from arousals or movement, the EEG signal was high-pass filtered at 150 Hz and the root mean square (RMS) signal calculated based on a window length of 100 ms. Again, an individual threshold was determined by the median + 4 * interquartile range of the rms signal and time points were marked as high frequency burst, when the RMS signal exceeded the corresponding threshold for at least 100 ms. Automated detection was followed by a visual inspection and all detected artifact samples were then padded by ± 250 ms. Furthermore, artifact-free intervals shorter than 3 s were also marked as artifacts.

#### Offline detection of discrete spindle and ripple events

Discrete spindles and ripples were detected during artifact-free NREM sleep using offline algorithms [14]. For spindle detection, the NC- and HIPP-signals were band-pass filtered at 12–16 Hz and the root mean square signal (RMS) was calculated based on a 200-ms windows followed by an additional smoothing with the same window length. A spindle event was identified whenever the smoothed RMS-signal exceed a threshold, defined by the mean plus 1.25 times the standard deviation of the RMS-signal across all NREM data points, for at least 0.4 s but not longer than 3 s. Importantly, time points exceeding an upper threshold determined by the mean RMS-signal plus 5 times its the standard deviation were excluded. The upward and downward threshold crossings represent the onset and end of a spindle event.

Detection of discrete ripple events in the hippocampal depth recordings followed the same procedure, except that the EEG signal was band-pass filtered from 80 to 120 Hz and both RMS calculation and smoothing were based on 20-ms windows. Detection and upper cut-off threshold were defined by the mean of the RMS-signal plus 2.5 or 9 times the standard deviation, respectively. Potential ripple events with a duration shorter than 38 ms (corresponding to 3 cycles at 80 Hz) or longer than 100 ms were rejected. Additionally, all ripple events were required to exhibit a minimum of three cycles in the raw EEG signal. For subsequent analysis all ripple events were segmented into 2-s epochs centred on their positive peak.

Finally, to ensure that both spindle and ripple events were not caused by spurious broadband power increases but reflect discrete events within our frequency range of interest, we implemented a routine to discard false positives based on their frequency profile. To this end, we calculated a time-frequency representation time-locked to the maximum of each spindle or ripple event (spindles: frequencies from 9 to 19 Hz in 0.5-Hz steps, time window of ± 750 ms in 2 ms steps; ripples: frequencies from 65 to 135 Hz in 2-Hz steps, time window of ± 100 ms in 2 ms steps) and extracted the frequency profile by averaging along the time dimension from −0.5 to +0.5 s for spindle events or from −0.05 to +0.05 for ripple events. A spindle/ripple event was rejected as a false positive whenever the frequency profile did not exhibit a prominent peak, i.e. a decline in amplitude on both sides of at least 20% with respect to its maximum value (determined with the prominence output of the MATLAB function ‘findpeaks’), within the frequency range of interest, i.e., between 12 to 16 Hz or 80 to 120 Hz, respectively.

#### Surrogate ripple events

To statistically assess of the dynamics around sharp-wave ripples, we used carefully matched surrogate data in which no ripples were observed. One advantage of this procedure is that it avoids the arbitrary decision of what constitutes a proper pre-ripple baseline period. In particular, for each participant’s n observed ripple events, we derived n non-ripple events, i.e. artifact-free NREM epochs matching the duration of each individual event including an additional padding of 1.5 s before and after in which our ripple detection algorithm did not indicate the presence of a ripple. Furthermore, to ensure that signal properties are maximally matched between target events and surrogates, surrogate events were only drawn from a 10 min time window before and after the corresponding ripple event with probabilities modulated according to a normal distribution. Epochs assigned to surrogate events were discarded from subsequent iterations to exclude overlapping non-events. This procedure was repeated 100 times. These 100 sets of matched surrogates were collapsed for power comparisons as shown in Figure 2 and used to build a surrogate distribution against which to compare the empirically observed connectivity metrics (Figures 3, S2-S4).

Visualisation of spindle and sharp-wave ripples waveform as shown in Figure 1 was performed by averaging the EEG signals time-locked to the minimum spindle trough or maximum ripple peak segmented into ± 1 s and ± 0.5 s epochs, respectively.

#### Time-frequency representations

Ripple-locked TFRs were calculated on the epoched ripple-events across all patients using Morlet wavelets for frequencies from 0.5 to 20 Hz with a 0.5 Hz resolution in 20 ms steps. For frequencies ≥5 Hz, the number of cycles was set adaptively to half of the corresponding frequency (or rounded up to the next integer value) but at least 5 cycles, resulting in time windows of approximately 500 ms. For frequencies below 5 Hz, i.e. 0.5 to 4.5 Hz, cycle numbers were reduced to values ranging from 1 to 4, reducing the window size and thereby increasing availability of artefact-free segments. TFRs on surrogate data were calculated with an identical procedure.

For statistical analysis of power changes, we first averaged across all 100 surrogate sets and then used a two-tailed paired-samples t tests to test for significant differences between the ripple and surrogate distributions. To correct for multiple comparisons, a cluster-based permutation procedure was applied as implemented in FieldTrip [49], using a cluster threshold of P < 0.05 and a final threshold for significance of P < 0.05.

#### Coherence

To assess the degree of synchronous activity between the neocortex and hippocampus around SW-Rs, time-resolved coherence from −1 to +1 centred on ripple events was calculated for frequencies from 0.5 to 20 Hz using cross-spectral densities from complex time-frequency representations (ft_connectivityanalysis function with method-parameter = ‘coh’ in FieldTrip). Statistical analysis was initially restricted to the spindle range (frequencies: 12 to 16 Hz, time: −0.25 to +0.25 s). To this end, the empirical value for coherence around ripples was obtained by averaging across our window of interest and tested for significance by means of a z-statistics using the mean and standard deviation of the corresponding coherence from the 100 surrogate datasets:

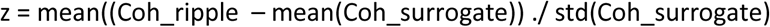

The z-values were than transformed into P-values, with a significance threshold of z-values greater or smaller than +/-1.96. Furthermore, to illustrate the temporal dynamics of coherence in the spindle range, we first averaged the resulting coherence representation across the frequency dimension from 12 to 16 Hz to obtain a coherence time series and then performed a z-transformation with the corresponding time series from the surrogate data.

#### Partial directed coherence

To examine directionality in the cortico-hippocampal communication we calculated partial directed coherence (PDC, ft_connectivityanalysis function with method-parameter = ‘pdc’ in Fieldtrip), which extends the concept of coherence of mutual synchrony by a decomposition into direct driving influence between the regions of interest, i.e., NC driving HIPP and vice versa [29]. In order to resolve directionality in time, we computed PDC on complex power spectra obtained on 512-ms long intervals shifted from −1 to +1 s around detected ripple events in 20-ms steps. Subsequently, statistical analyses focused on mean PDC values obtained by averaging across 12-16 Hz and a time interval from −0.25 to +0.25 s. Again, to illustrate the temporal evolution of PDC in the spindle range, we first extracted the corresponding time series by averaging from 12 to 16 Hz and then calculated a z-score with respect to the surrogate data. Finally, we subtracted the resulting z-scores for the two directions, i.e. NC driving HIPP minus HIPP driving NC. Thus, positive values indicate a driving influence of neocortical on hippocampal spindle activity and vice versa for negative values. Given the ambiguity of any directional influence in the absence of functional coupling, we restrict display and interpretation of PDC to time/frequency bins exhibiting positive spectral coherence in the preceding analysis.

#### Amplitude- and phase-based connectivity

Our examination of mutual and directed connectivity based on spectral and partial directed coherence has the advantage that both rely on the same mathematical framework and thus allow for internally consistent statistical assessments (i.e., PDC informing on the directionality of the observed spectral coherence). Nonetheless, we asked to what extend hippocampal-neocortical communication is mediated by cross-regional amplitude or phase-relationships. To disentangle this, we additionally calculated the power-power correlation [27] and phase locking value (PLV, [28]) between HIPP and NC during ripples and surrogate events.

It is important to note that connectivity measures are prone to volume conduction (despite long distances between regions of interest) or spurious coupling effects introduced by a common reference. To mitigate this concern we removed signal components exhibiting identical instantaneous phase by orthogonalizing our data before calculating the power-power correlation between HIPP and NC [27]. More specifically, for each ripple/surrogate event, the complex TFR for NC was orthogonalized with respected to HIPP (NC ⊥ HIPP) and HIPP orthogonalized with respect to NC (HIPP ⊥ NC) prior to computing the absolute power. Next, for each time-frequency point we calculated the correlation between NC and (HIPP ⊥ NC) and between (NC ⊥ HIPP) and HIPP across all trials and averaged both resulting correlation maps.

Finally, we examined phase-locking between HIPP and NC [28]. Based on complex TFRs for frequencies from 0.5 to 20 Hz and between −1 s to +1 s around ripple/surrogate events, the PLV (ft_connectivityanalysis function with method-parameter = ‘plv’ in Fieldtrip) extracts the phase information and evaluates the variation in phase-difference between HIPP and NC for a given frequency and time point across all ripple and surrogate events, respectively.

#### Peri-event histograms of spindle onsets

Finally, given our findings on the directional influence between NC- and HIPP-spindles we inspected the timing of spindle onsets within both regions of interest. To this end, we created peri-event histograms (bin size = 50 ms) of the onsets of discrete spindle events time-locked to ripple and surrogate events occurring between −0.5 to +0.5 s. The resulting histograms were normalised by the total number of detected spindle onsets (multiplied by 100) and smoothed by a three-point moving average. Identical to our previous analyses, statistical comparison with the surrogate data was performed by calculating a z-value with respect to the mean and standard deviation of the surrogate distribution for each region and bin.

