## Supplementary Material for "Sleep spindles mediate hippocampal-neocortical coupling during sharp-wave ripples"

**Supplemental items**

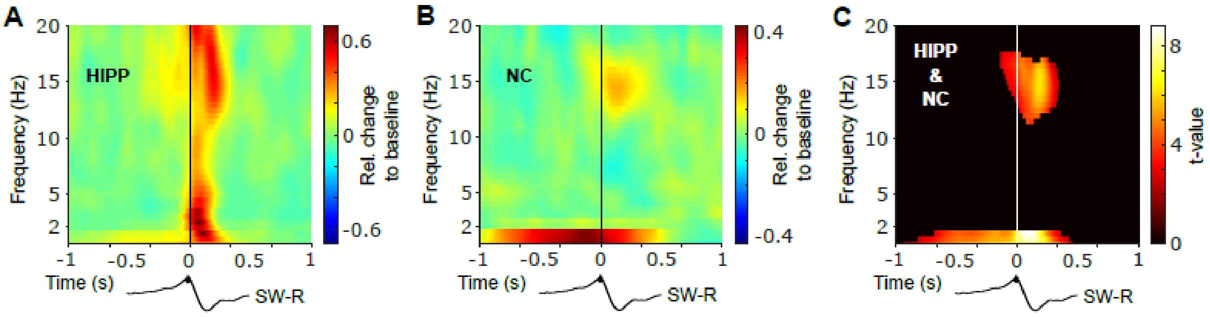


**Figure S1. SW-R locked TFRs relative to pre-event baseline. Related to Figure 2.** (A) SW-R locked TFRs in HIPP (left) and NC (right). Color code reflects relative change in power with respect to a baseline from -2 to -1.5. (B) Significance mask derived from the overlap of significant clusters between NC and HIPP. Color represents the mean t-value from the corresponding statistical masks. Black traces below illustrate the timing relative to HIPP SW-Rs.

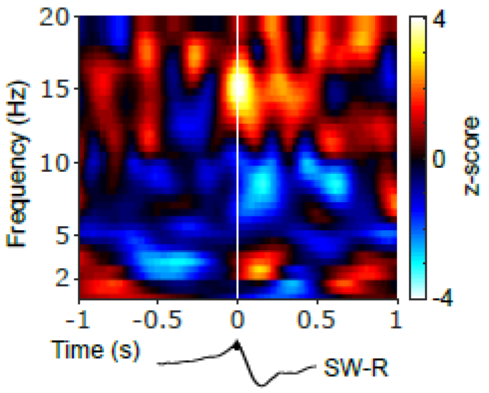


**Figure S2. Cortico-hippocampal spectral coherence around SW-Rs. Related to Figure 3.** Statistical map (z-scores) contrasting ripple-locked and surrogate coherence between NC and HIPP from 0.5 to 20 Hz and -1 to 1 s around SW-Rs.. Black traces below illustrate the timing relative to HIPP SW-Rs.

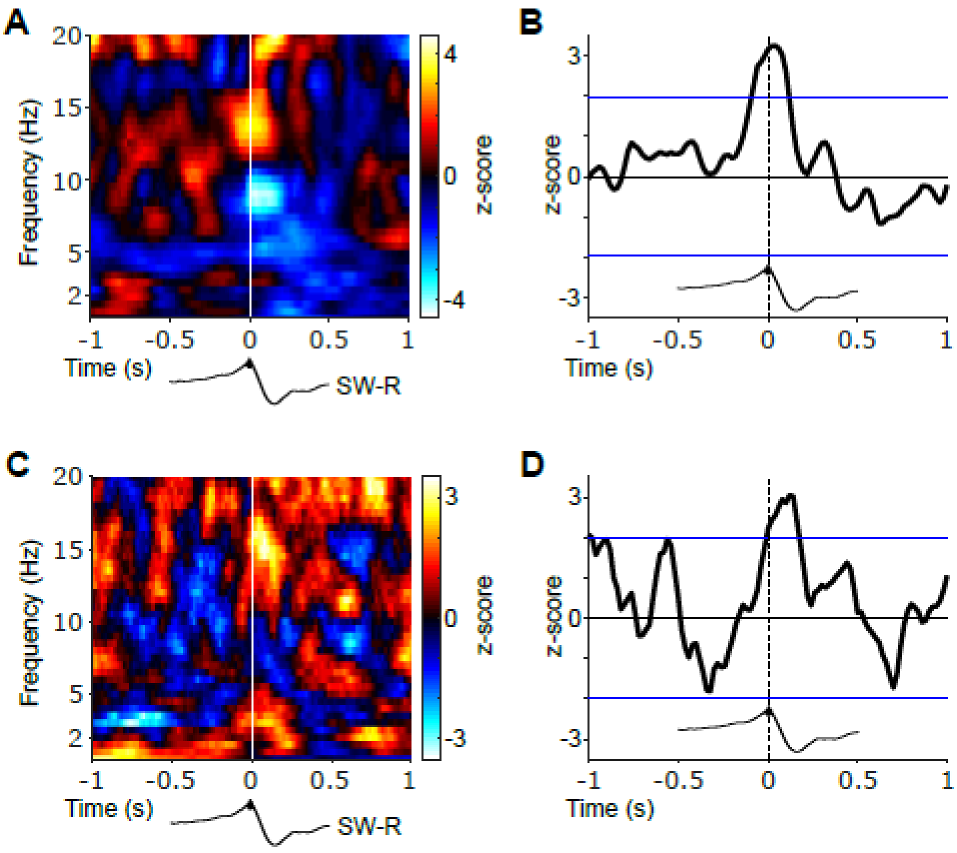


**Figure S3. Amplitude- and phase-based cortico-hippocampal connectivity. Related to Figure 3.** (A) Statistical map (z-scores) contrasting ripple-locked and surrogate connectivity based on the orthogonalized power correlation. (B) Time-resolved NC-HIPP power correlation for the 12-16 Hz spindle range were transformed into a z-score with respect to the surrogate events. Blue lines indicate standard significance thresholds (z = 1.96). Time 0 denotes HIPP SW-Rs. (B) Statistical map (z-scores) contrasting ripple-locked and surrogate connectivity assessed via phase-locked value (PLV). (D) Time-resolved NC-HIPP PLV for the 12-16 Hz spindle range transformed into a z-score with respect to the surrogate events. Blue lines indicate standard significance thresholds (z = 1.96). Time 0 denotes HIPP SW-Rs.

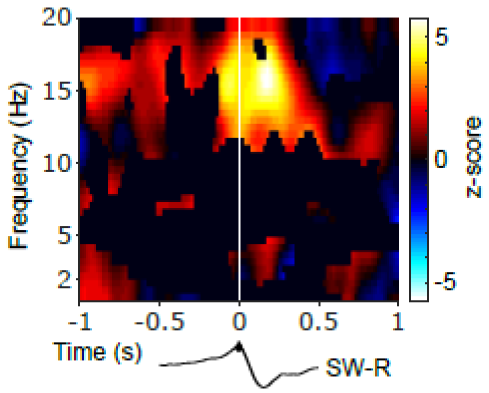


**Figure S4. Directed cortico-hippocampal connectivity around SW-Rs.** Statistical map (z-score), thresholded based on positive spectral coherence (Figure S2), depicting the difference in PDC between NC->HIPP and HIPP->NC time-locked to ripples in comparison to surrogates. Hot colors indicate a top-down influence from NC to HIPP, whereas cold colors indicate a reversed directional influence, i.e. HIPP driving NC. Black traces below illustrate the timing relative to HIPP SW-Rs.
